# The role of the *C. albicans* transcriptional repressor *NRG1* during filamentation and disseminated candidiasis is strain-dependent

**DOI:** 10.1101/2023.12.15.571891

**Authors:** Rohan S. Wakade, Melanie Wellington, Damian J. Krysan

## Abstract

*Candida albicans* is one of the most common causes of superficial and invasive fungal disease in humans. Its ability to cause disease has been closely linked to its ability to undergo a morphological transition from budding yeast to filamentous forms (hyphae and pseudohyphae). The ability of *C. albicans* strains isolated from patients to undergo filamentation varies significantly. In addition, the filamentation phenotypes of mutants involving transcription factors that positively regulate hyphal morphogenesis can also vary from strain to strain. Here, we characterized the virulence, in vitro and in vivo filamentation, and in vitro and in vivo hypha-associated gene expression profiles of four poorly filamenting *C. albicans* isolates and their corresponding deletion mutants of the repressor of filamentation *NRG1*. The two most virulent strains, 57055 and 78048, show robust in vivo filamentation while remaining predominately yeast phase exposed to RPMI+10% bovine calf serum at 37°C; the two low virulence strains (94015 and 78042) do not filament well under either condition. Deletion of *NRG1* increases hyphae formation in the SC5314 derivative SN250 but only pseudohyphae are formed in the clinical isolates in vivo. Deletion of *NRG1* modestly increased the virulence of 78042 which was accompanied by increased expression of hyphae-associated genes without an increase in filamentation. Strikingly, deletion of *NRG1* in 78048 reduced filamentation, expression of candidalysin (*ECE1*) and virulence in vivo without dramatically altering establishment of infection. Thus, the function of *NRG1* varies significantly within this set of *C. albicans* isolates and can actually suppress filamentation in vivo.

**Importance:** Clinical isolates of the human fungal pathogen *Candida albicans* show significant variation in their ability to undergo in vitro filamentation and in the function of well-characterized transcriptional regulators of filamentation. Here, we show that Nrg1, a key repressor of filamentation and filament specific gene expression in standard reference strains, has strain dependent functions, particularly during infection. Most strikingly, loss of *NRG1* function can reduce filamentation, hypha-specific gene expression such as the toxin candidalysin, and virulence in some strains. Our data emphasize that the functions of seemingly fundamental and well-conserved transcriptional regulators such as Nrg1 are contextual with respect to both environment and genetic background.

## Introduction

*Candida albicans* is a component of the human mycobiome and one of the most common causes of human fungal infections in both immunocompetent and immunocompromised people (1). The ability of *C. albicans* to undergo morphological transitions between round, budding yeast forms and the filamentous morphologies of pseudohyphae and hyphae has been strongly correlated with the ability of *C. albicans* to disease (2). Accordingly, this virulence trait has been studied extensively, leading to the identification of a large number of genes that affect *C. albicans* filamentous morphogenesis (3). The majority of these genes identified by experimentation with a single strain background, SC5314, either directly or through work with its auxotrophic derivatives.

In recent years, interest in the characterization of clinical isolates of *C. albicans* has increased (4, 5, 6). Through this work, varying degrees of phenotypic heterogeneity have been identified particularly with respect to in vitro filamentation and biofilm formation. Studies of a set of 20 clinical isolates predominantly from bloodstream infections (7) have found that in vitro filamentation does not correlate well with virulence phenotypes in mouse models of invasive disease (4). A large study of commensal isolates of the GI tract also found a large amount of in vitro phenotypic diversity whereas virulence phenotypes in an invertebrate model were quite consistent regardless of in vitro phenotype (8). Furthermore, in vitro studies of the function of transcription factors have also revealed that the strain background can have profound effects on the function of factors such as *BCR1* and *NRG1* (6, 9). Here, we describe our investigation of the effect of strain background on the role of the transcriptional repressor Nrg1 during in vitro filamentation, in vivo filamentation and disseminated candidiasis in a mouse model.

*NRG1* is a repressor of both filamentation and hypha-associated gene expression in *C. albicans* and its deletion in SC5314 derivatives leads to constitutive pseudohyphae formation under non-inducing conditions in vitro (10, 11). At the initiation of normal in vitro filamentation, *NRG1* expression is reduced and its gene product, Nrg1, is degraded (12). This inhibition of Nrg1 function is correlated with the expression of hyphae-specific genes, many of which appear to be direct Nrg1 binding targets (11). In contrast, constitutive expression of *NRG1* blocks in vitro and in vivo filamentation and prevents the development of disease, but not organ infection, in a mouse model of disseminated infection (13). In contrast, reduced expression of *NRG1* leads to increases in in vitro and in vivo filamentation and disease in the mouse model.

Because of its profound effects on filamentation and virulence in the SC5314 background, we were interested to explore the effect of strain background on *NRG1* function. To do so, we deleted *NRG1* in four poorly filamenting clinical isolates and examined the effect of those mutations on in vitro and in vivo filamentation and gene expression as well as on virulence in the mouse disseminated candidiasis model. Although *nrg1*ΔΔ mutants have relatively similar phenotypes and effects on gene expression in vitro across these strains, there are profound strain-based differences during mammalian infection.

## Results

### *NRG1* deletion mutants in poorly filamenting clinical isolates establish infection but their effect on virulence is isolate-dependent

We selected four *C. albicans* clinical isolates (94015, 57055, 78048, and 78042) that showed very low levels of filamentation after induction with RPMI medium at 37°C as reported by Hirakawa et al (4). Relative to SC5315 and its derivatives, all four strains were also shown previously to have reduced virulence in the mouse model of disseminated candidiasis (7). As expected, *NRG1* deletion mutants in SN250 form pseudohyphae in non-filament inducing conditions (Fig. 1A&B). In contrast, the *nrg1*ΔΔ mutants derived from the four clinical isolates are predominately yeast phase in rich medium cultures at 30°C (Fig. 1A&B) and form approximately 20% pseudohyphae.

**Figure 1.**
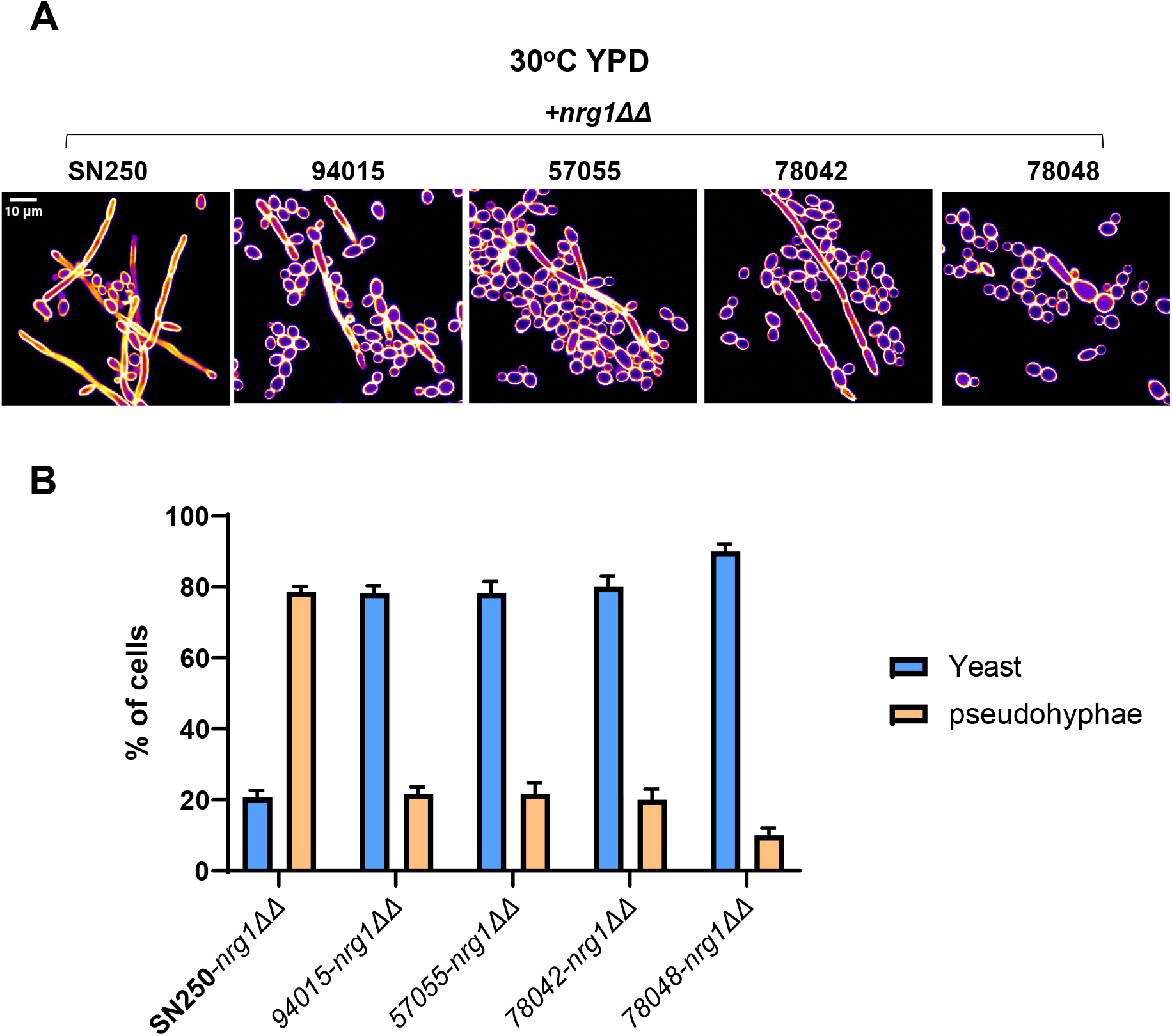
Morphology of *NRG1* deletion mutants in non-hyphae inducing conditions. **A.** Representative images of *nrg1*ΔΔ mutants in the indicated strain background after growth in Yeast Peptone Dextrose medium in 30°C and Calcofluor White staining. **B.** Quantitation the yeast and pseudohyphal forms in the *nrg1*ΔΔ mutants in the indicated strain backgrounds. The bars indicate mean of three independent experiments with at least 100 cells counted in multiple microscopy fields. Error bars are standard deviation.

We inoculated the four clinical isolates and their corresponding *nrg1ΔΔ* mutants in outbred CD-1 mice and determined the kidney fungal burden at post-infection day 3. No mortality was observed prior to harvest and all four parental strains established infections with between 10^5^ and 10^6^ colony forming units (CFU) per mL of kidney homogenate (Fig. 2A-D). The corresponding *nrg1*ΔΔ mutants also established infections. Mice infected with *nrg1*ΔΔ mutants in the 94015 (Fig. 2A), 57055 (Fig. 2B), and 78042 (Fig. 2C) backgrounds showed no statistically significant difference in kidney burden relative to the parental strains (Student’s t test of the log^10^ transformed data, p > 0.05), whereas mice infected with the *nrg1*ΔΔ-78048 strain showed reduced fungal burden relative to the parental stain (Fig. 2D). We did not test the SN250-derived *nrg1*ΔΔ mutant because the extent of pseudohyphae formation precludes accurate inoculum quantitation.

**Figure 2.**
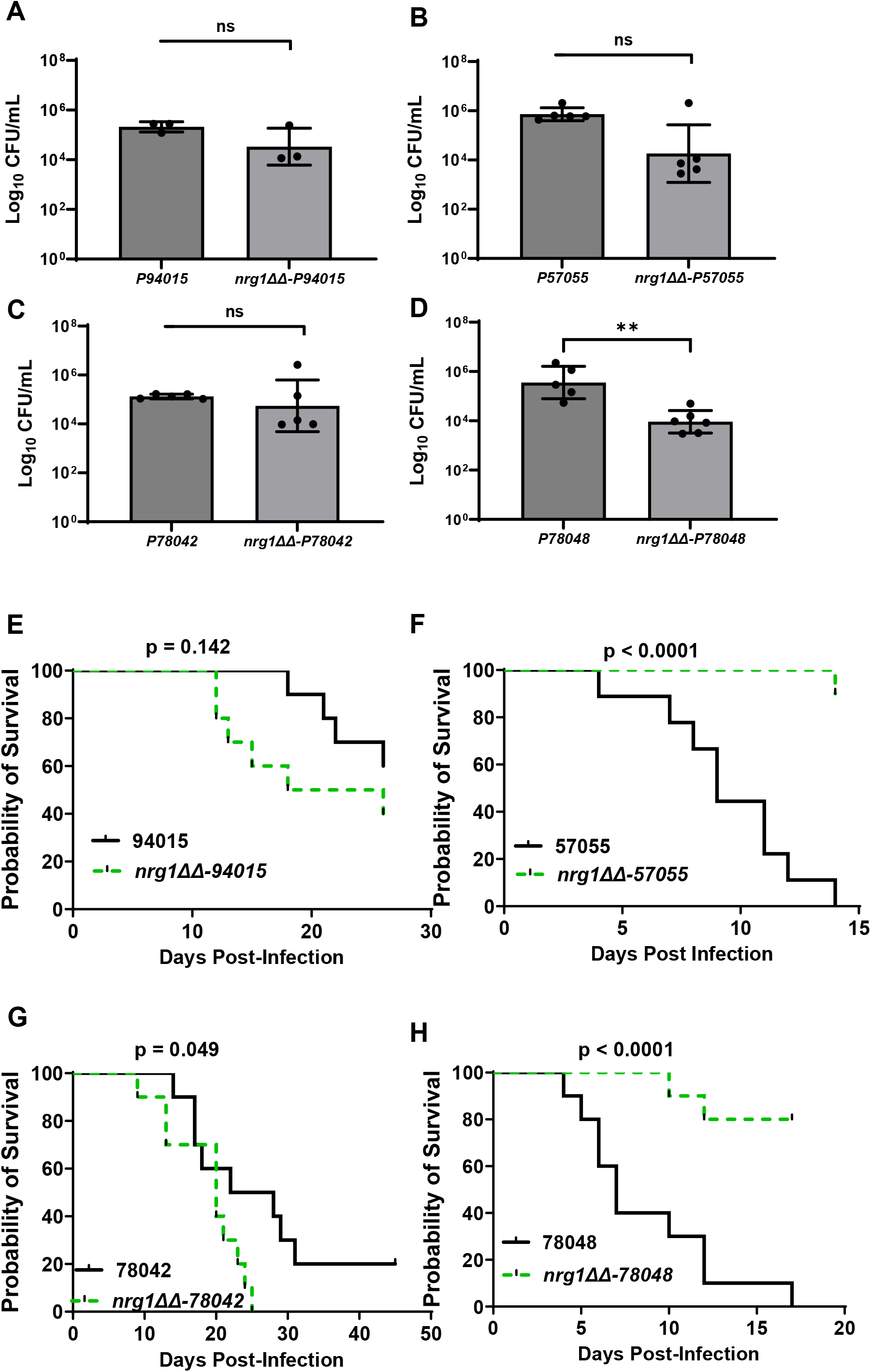
A *nrg1*ΔΔ mutation has strain dependent effects on infectivity and virulence. Kidney fungal burden 3 days post-infection for the indicated clinical strains (**A-D**) by tail-vein and its corresponding *nrg1*ΔΔ mutant (n = 5 mice per group) by tail vein injection. Bars indicate mean of log^10^ fungal burden (CFU/mL) with individual mice shown as points and error bars indicating standard deviation. Differences between groups were analyzed by Student’s t test (NS: p> 0.05 and * p< 0.05). Survival curves for mice infected with the indicated clinical strains (**A-D**) and its corresponding *nrg1*ΔΔ mutant (n = 10 mice per group) by tail vein injection. The curves indicate survival as defined as time to moribundity. P values are for Log Rank (Mantel-Cox) analysis with statistical significance defined as p <0.05.

Having confirmed that the parental strains and their corresponding *nrg1*ΔΔ mutants established infection in the mouse model of disseminated candidiasis, we next compared their effect on virulence (Fig. 2E-G). Importantly, all clinical isolates all caused some level of disease with 57055 and 78042 causing moribundity (median survival time: 57055 (9D); and 78048 (7D)) comparable to previously reported data for SN250 in the same mouse strain background (median survival time: SN250 (6D); ref. 14). The strain lacking a functional allele of *EFG1* (4), 94015, was the least virulent with only 40% moribundity 4 weeks post infection (Fig. 2E; median survival time; undefinable); 78042 also showed low virulence with 80% moribundity at 4 weeks that did not increase further over an additional 3 weeks (Fig. 2F). Despite the poor filamentation of these strains in vitro, they were able to cause disease but, as previously described (7), to differing extent.

Based on the behavior of conditionally regulated *NRG1* derivatives SC5314 (13), we hypothesized that the *nrg1*ΔΔ mutants might increase the virulence of the poorly filamenting strains by promoting filamentation or hyphae-associated gene expression. This hypothesis appears to be partially correct. Deletion of *NRG1* in the least virulent 94015 strain did not affect virulence in a statistically significant manner, although there was a trend toward increased virulence (Fig. 2E). We examined the fungal burden of four surviving animals from both groups and found that the 94015 strain had been cleared in 3/4 animals while all four of the surviving animals infected with *nrg1*ΔΔ-94015 had robust kidney fungal burden (Fig. S1A). Consistent with our initial hypothesis, the *nrg1*ΔΔ-78042 strain had a modest increase in virulence relative to the parental strain (Fig. 2G).

In contrast, deletion of *NRG1* in the two more virulent strains, 57055 and 78048, reduced virulence substantially in both strains (Fig. 2F&H). The *NRG1* mutants of both strains caused reduced fungal burden at day 3 with the difference between the *nrg1*ΔΔ-78048 and 78048 being statistically significant (Fig. 2B&D). Thus, it is possible that the reduced initial fungal burden of these mutants contributes to their lower virulence relative to the corresponding parental strains. Interestingly, the fungal burden of the kidneys of surviving mice infected with either *nrg1*ΔΔ-57055 or *nrg1*ΔΔ-78048 was 10^5^ and 10^6^ CFU/mL, respectively (Fig. S1B&C). The reduced virulence of these two strains, therefore, is not due to clearance of the fungus. This indicates that loss of *NRG1* function does not drive increased virulence in all strains.

### Deletion of *NRG1* in poorly filamenting clinical isolates increases pseudohyphae formation in vitro

Hirakawa et al. had found that none of the four clinical isolates formed significant filaments when induced with RPMI tissue culture medium for 6 hr at 37°C (4). We tested their ability to filament in RPMI+10% bovine calf serum (BCS) at 37°C for 4hr to determine if the addition of serum increased their filamentation. For all strains, yeast remained the predominant morphology under these conditions (Fig. 3A&B). All strains formed some filamentous forms with pseudohyphae outnumbering true hyphae and 94015 forming very low numbers of pseudohyphae (Fig. 3A&B). Thus, the addition of serum did not greatly affect the in vitro phenotypes and did not provide insights into differences in virulence between the different strains.

**Figure 3.**
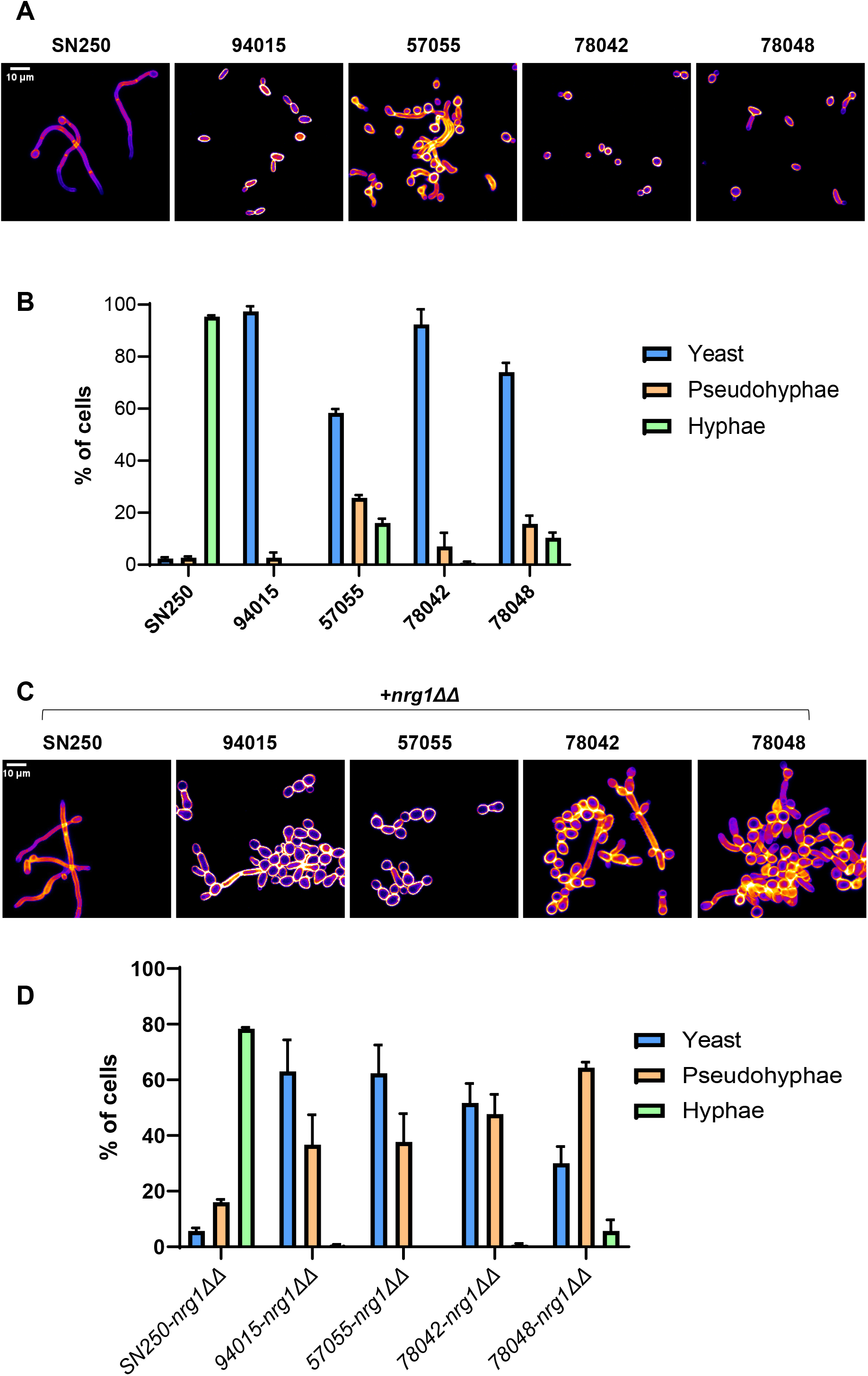
Clinical strains show low rates of filamentation in vitro and deletion of *NRG1* leads to increased pseudohyphae. **A**. Representative images of indicated strains after incubation in RPMI+10% bovine calf serum for 4hr at 37°C and Calcofluor White staining. **B.** Quantitation the yeast, pseudohyphal, and hyphal forms in the *nrg1*ΔΔ mutants in the indicated strain backgrounds. The bars indicate mean of three independent experiments with at least 100 cells counted in multiple microscopy fields. Error bars are standard deviation. **C**. Representative images of *nrg1*ΔΔ mutants in the indicated strain background after incubation in RPMI+10% bovine calf serum for 4hr at 37°C and Calcofluor White staining. **D**. Quantitation the yeast, pseudohyphal, and hyphal forms in the *nrg1*ΔΔ mutants in the indicated strain backgrounds. The bars indicate mean of three independent experiments with at least 100 cells counted in multiple microscopy fields. Error bars are standard deviation.

Exposure of the *nrg1*ΔΔ mutants to filament-inducing conditions led to pseudohyphae formation in all strain backgrounds (Fig. 3C&D). The 94015-*nrg1*ΔΔ and 57055-*nrg1*ΔΔ mutants formed ∼2:1 ratio of yeast to pseudohyphae with no hyphae observed. The 78042-*nrg1*ΔΔ mutant formed a 1:1 ratio of yeast to pseudohyphae while the 78048-*nrg1*ΔΔ mutant formed predominantly pseudohyphae with a small number of true hyphae observable. The predominance of pseudohyphae in these *nrg1*ΔΔ mutants is distinct from the SN250-*nrg1*ΔΔ mutant which forms 80% hyphae (Fig. 3D). Although deletion of the repressor of filamentation *NRG1* in these low filamenting clinical isolates increases filamentation relative to their parental strains under inducing conditions, the strains form pseudohyphae instead of true hyphae, indicating that their inability to form true hyphae is unlikely to be due to a failure to inhibit Nrg1 repression under filament-inducing conditions.

We also examined the filamentation phenotypes of the clinical isolates and their corresponding *NRG1* deletion mutants on solid media (RPMI and RPMI+10%BCS). The only two parental strains that showed significant filamentation at either 30°C or 37°C were SN250 and 78048 (Fig. S2A&B, respectively). In all strains, the *NRG1* deletion mutants formed highly wrinkled colonies at both 30°C and 37°C in both serum-free and serum-containing medium (Fig. S2B). Peripheral invasion was evident at 30°C for all mutants in RPMI (but not RPMI+10%BCS) and was robust in the SN250 and 78048 derivatives; however, this invasion was lost in all but the SN250- and 78048-*nrg1*ΔΔ mutants at 37°C. Thus, during filamentation on solid medium, the colony morphology phenotypes of *nrg1*ΔΔ mutants are reasonably consistent across this set of four strain backgrounds.

Taken together, the in vitro filamentation phenotypes of both the parental and the *nrg1*ΔΔ mutants are fairly consistent across the four clinical isolates. Thus, it is not possible to explain the differences in the virulence of either the parental or *nrg1*ΔΔ mutants using these phenotypes. It is, however, very clear that, in contrast to *nrg1*ΔΔ-SN250, the clinical isolate-derived *nrg1*ΔΔ mutants form pseudohyphae rather than hyphae under inducing conditions. It appears that Nrg1 suppresses filamentation in these clinical isolates but that the formation of hyphae is dependent upon additional factors that are absent or reduced in the poorly filamenting clinical isolates relative to SN250.

### In vivo filamentation phenotypes of the clinical isolates and their *nrg1*ΔΔ mutants are strain dependent

In agreement with its inability to filament in vitro, we previously reported that 94015 also failed to undergo filamentation in vivo (15). In contrast, 50755 forms filaments in vivo to a similar extent as SN250 and SC5314 despite having reduced filamentation under a variety of in vitro conditions (Fig. 3A&B, ref. 9). Therefore, poor in vivo filamentation of 94015 correlates with its low virulence and robust in vivo filamentation correlates with the ability of 57055 to cause disease. To further test this correlation, we examined the in vivo filamentation phenotypes of 78042 and 78048. Consistent with the phenotypes for 50755 and 94015, the low virulence 78042 strain forms essentially no filaments in vivo (Fig. 4A) while the virulent 78048 forms filaments (Fig. 4B, 68%) at a rate we previously observed for 50755 and SN250 (15). Thus, the in vivo filamentation phenotypes of these strains correlate well with their virulence while the in vitro filamentation phenotypes do not.

**Figure 4.**
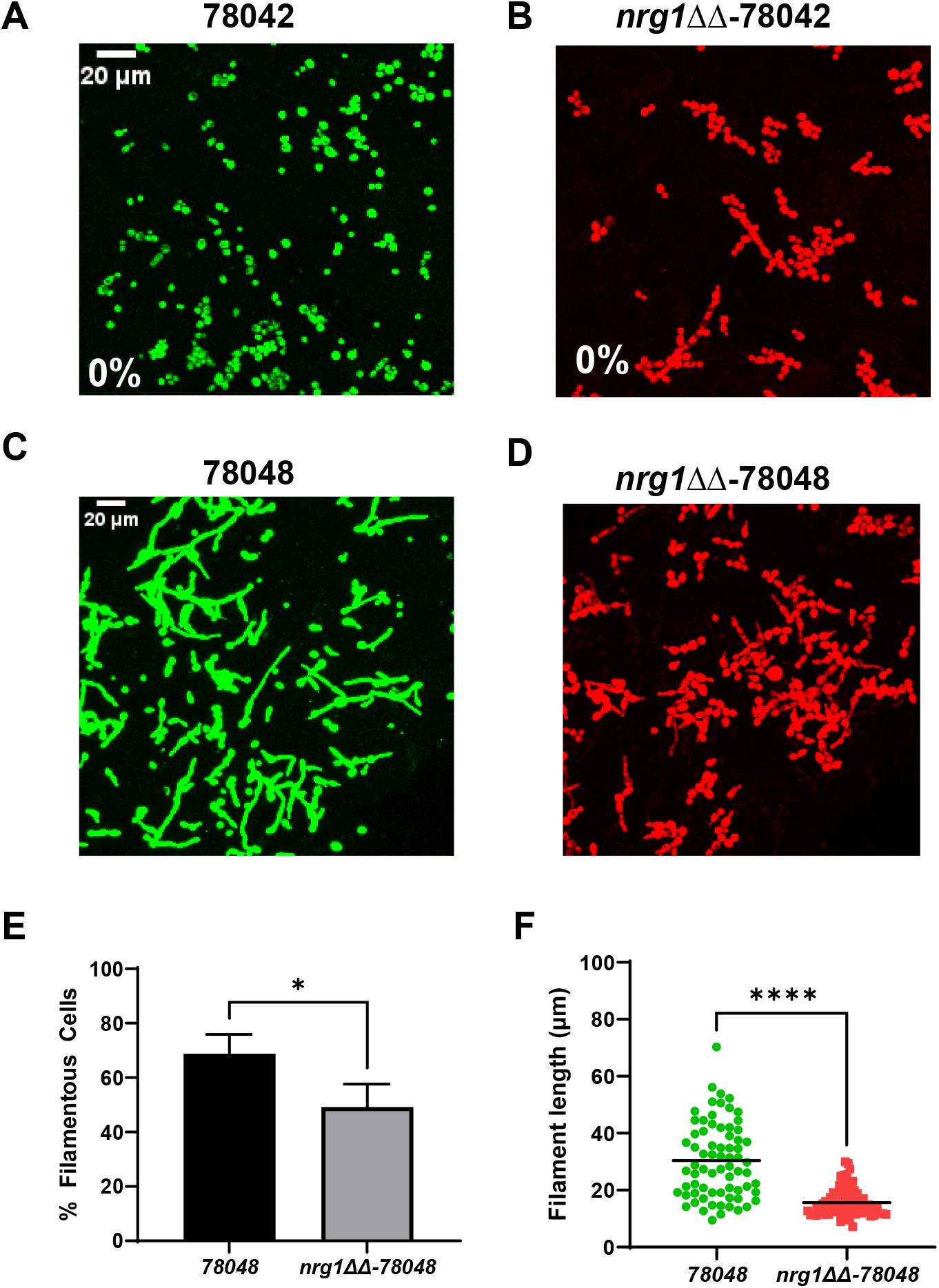
Effect of *NRG1* deletion on in vivo filamentation in clinical strains 78042 and 78048. Representative images for mNEON-78042 (**A**), iRFP-*nrg1*ΔΔ-78042 (**B**), mNEON-78048 (**C**), and iRFP-*nrg1*ΔΔ-78048 (**D**) taken 24hr post-infection of ear tissue as described in materials and methods. For the 78042 strains no filamentous cells were observed as indicated by the 0% in the image. **E**. Quantitation of % filamentous cells for the 78048 strains using the scoring criteria described in materials and methods. The bars indicate at least two independent replicates with standard deviation indicated by error bars. * indicates p value <0.05 by Student’s t test. **F**. Filament length for the 78048 strains dots indicate individual filaments with data pooled from two independent infections. **** indicates p <0.00001 by Mann-Whitney U test.

We focused our in vivo filamentation analysis of the *nrg1*ΔΔ mutants on 78042, a poorly filamenting strain in vivo, and 78048, a robustly filamenting strain in vivo. Despite showing increased virulence relative to its parental strain, the *nrg1*ΔΔ-78042 strain remained in the yeast morphology in vivo with no clearly filamentous forms identified (Fig. 4C). Strikingly, the *nrg1*ΔΔ-78048 strain showed a statistically significant reduction in the proportion of filaments and a significant shortening of the filament lengths relative to its parental strain (Fig. 4D, E&F). Although it’s not possible to conclusively distinguish hyphae and pseudohyphae with this in vivo imaging assay, pseudohyphae have shorter filament lengths. Thus, the *nrg1*ΔΔ-78048 mutant may be forming more pseudohyphae relative to the parental strain or may be forming shorter hyphae, a feature also correlated with reduced virulence (15). Therefore, the reduced virulence observed for the *nrg1*ΔΔ-78048 mutant may be due in part to reduced filamentation. On the other hand, the increased virulence of the *nrg1*ΔΔ-78042 strain cannot be attributed to increased filamentation.

### In vitro expression of hyphae-associated genes is reduced in poorly filamenting clinical isolates relative to SN250 and increases in their *NRG1* deletion mutants

A well-characterized set of hypha-associated genes are induced during filamentation and some directly contribute to virulence (e.g., *ECE1*). Thus, changes in the expression of hyphae-associated genes could contribute to differences in virulence for the clinical isolates and their respective *nrg1*ΔΔ mutants. To explore this further, we first characterized the expression profile of the clinical isolates by Nanostring under in vitro induction and compared the profiles to the strongly filamenting, SC5314-derivative SN250. The probe set contains 186 environmentally responsive genes including 57 hyphae-associated transcripts (see Table S1 for complete gene list). Volcano plots showing the fold-change in expression relative to SN250 for each clinical strain are shown in Fig. 5A; the number of differentially expressed genes (DEGs; log^2^ FC ±1 with FDR 0.1, Benjamini-Hochberg procedure) for each clinical isolate was: 94015 (42 up; 63 down): 57055 (18 up/35 down); 78042 (17 up/42 down); and 78048 (20 up/37 down).

**Figure 5.**
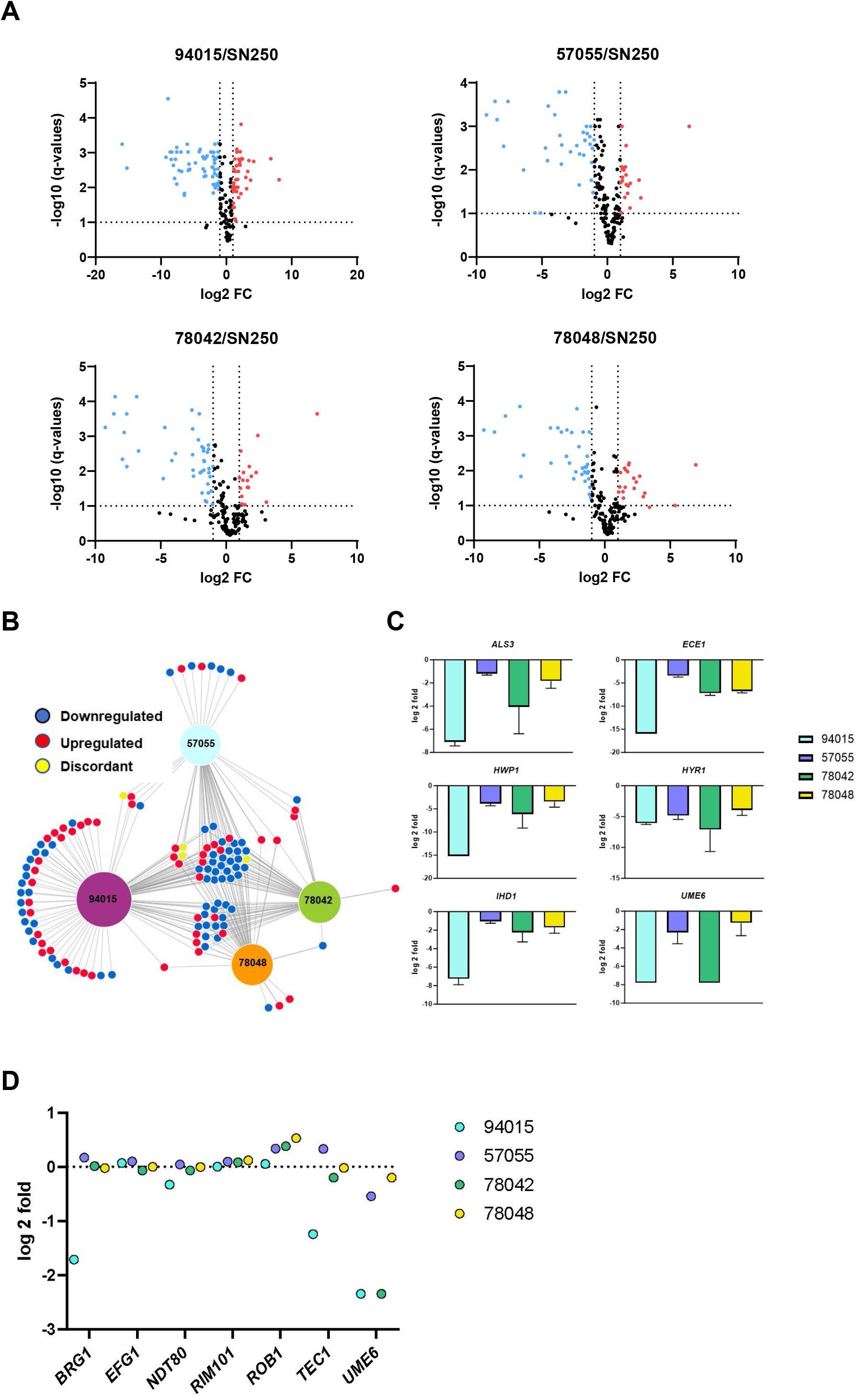
Poor filamenting clinical isolates show reduced expression of hyphae-specific genes in vitro. **A**. Volcano plots of gene expression of a set of 185 genes as characterized by Nanostring nCounter. The expression was normalized to the strongly filamenting reference strain SN250. The horizontal bar indicates FDR 0.1 (Benjamini-Hochberg) and the vertical line indicates log^2^ = 1 of the fold change which are the cutoff values for the definition of differentially expressed genes. **B**. Network diagram showing differentially expressed genes that are unique or common between the different clinical strains. The size of the hubs for each clinical isolate is proportional to the number of differentially expressed genes relative to SN250. **C**. Expression of representative hypha-induced genes for the indicated strains normalized to expression in SN250. Bars indicate the mean fold change of three independent replicates and error bars indicate standard deviation. **D**. Log_2_ fold change for the expression of the indicated hyphae-associated transcription factors in the four clinical isolates normalized to SN250.

As shown in Fig. 5B, 94015 had the largest set uniquely downregulated genes (23). In contrast, the other three strains had very similar sets of downregulated genes. Twenty-three genes were downregulated in all four strains and included hyphae-associated genes; representative examples of these are shown in Fig. 5C. Finally, *YWP1* is expressed during yeast phase growth and is repressed during hyphal growth (16); all clinical strains showed increased expression of *YWP1* relative to SN250 (Table S1). Thus, the expression profile of the poorly filamenting strains is consistent with their in vitro filamentation phenotype.

### Lack of a functional *EFG1* allele provides a mechanism for the inability of 94015 to filament

(4). Therefore, one possible mechanism for the reduced filamentation of the other strains is reduced expression of the other transcription factors (TF) that are required for in in vitro filamentation relative to SN250. With the exception of 94015, the expression of *BRG1*, *EFG1*, *NDT80*, *RIM101*, *ROB1* and *TEC1* did not differ significantly from SN250 (Fig. 5D). Efg1 activates the expression of *BRG1* and *TEC1* during filamentation (15) and these TF genes were expressed at much lower levels in 94015. *UME6* is a TF required for the maintenance phase of filamentation as well as a marker of the filament expression program because it is not expressed at appreciable levels until filamentation is initiated (17). *UME6* expression is undetectable in yeast and remains so in 94015 as well as in 78042 during in vitro filamentation (Fig. 5D); consistent with this observation, these two strains form almost no hyphae or pseudohyphae. In contrast, *UME6* expression is expressed at levels near that of SN250 in 57055 and 78048, the two strains which form the highest levels of filaments in vitro; however, SN250 forms hyphae while 57055 and 78048 form pseudohyphae. This suggests that *UME6* expression is not sufficient to drive hyphae formation in these strains and that alteration in other aspects of the hyphal program are likely responsible for the lack of hyphae formation in 57055 and 78048.

We also generated Nanostring expression profiles for the *nrg1*ΔΔ deletion mutants during in vitro filament-induction (See Fig. S3A-D for volcano plots for each mutant compared to its parental strain; see Table SX for raw, processed and FC values along with FDR for each comparison). Although the clinical strain-derived *nrg1*ΔΔ mutants do not form true hyphae like the SN250-derived mutants, we expected they might express hyphae-associated genes at higher levels. The total number of DEGs in the *nrg1*ΔΔ deletion mutants of the clinical strains was: 49015 (XX; Y Up/ Z down); 57055 (XX; Y Up/ Z down); 78042 (XX; Y Up/ Z down); 78048 (XX; Y Up/ Z down).

Thirty-five genes are differentially expressed in a concordant manner in all four *nrg1*ΔΔ mutants (Fig. 6A). This set includes hypha-associated genes that are upregulated in all backgrounds as shown in Fig. 6B. We previously found that deletion of *NRG1* in an *efg1*ΔΔ mutant (SN250 background) restored expression of hypha-associated genes in vivo and in vitro (15). Consistent with the previous observations, hyphae-specific gene expression is increased *nrg1*ΔΔ-94015 (Fig. 6B) even though the strain does not form hyphae in vitro (Fig. 3C). The *nrg1*ΔΔ-94015 strain has the largest set of uniquely affected genes amongst the four strains with *nrg1*ΔΔ-50755 the strain has the next most. The set of differentially expressed in the *nrg1ΔΔ*-78042 and *nrg1ΔΔ*-78048 are almost completely shared with each other and the other two *NRG1* mutants (Fig. 6A). These data are consistent with results from the Mitchell lab indicating that the effect of transcriptional regulators on gene expression shows both similarities and distinctions across different strain background (5, 6).

**Figure 6.**
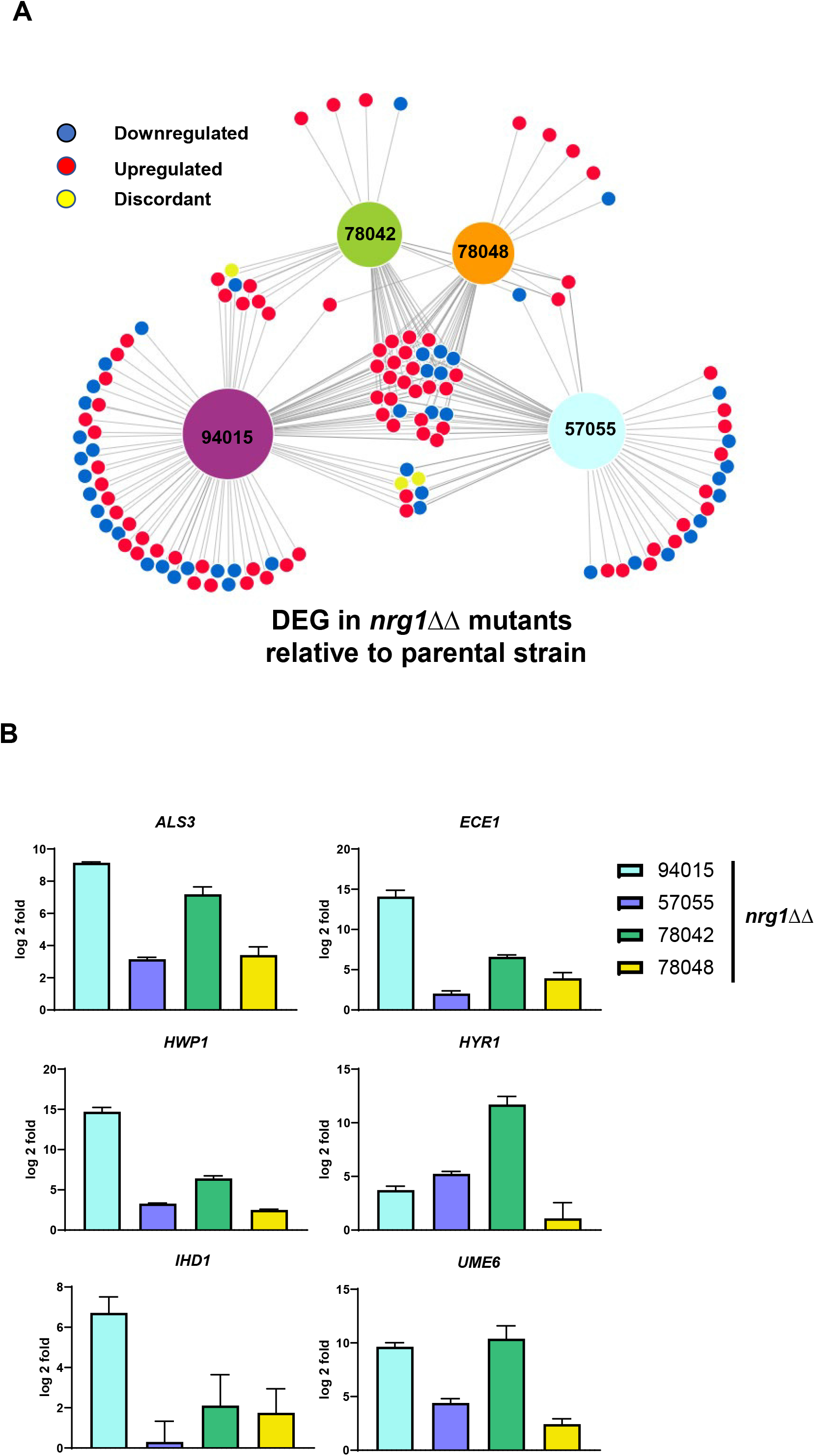
Deletion of *NRG1* increases expression of hypha-associated genes in the clinical isolate backgrounds in vitro. **A**. Network diagram showing differentially expressed genes in the *nrg1*ΔΔ mutants relative to the corresponding parental strains. The size of the hubs for each clinical isolate is proportional to the number of differentially expressed genes in the *nrg1*ΔΔ relative to its parent. **B**. Expression of representative hypha-induced genes for the *nrg1*ΔΔ mutant normalized to expression in its corresponding parental strain. Bars indicate the mean fold change of three independent replicates and error bars indicate standard deviation.

### Strain-dependent differential expression of hypha-associated in vivo correlates with the effect of *nrg1*ΔΔ mutation on virulence of 78042 and 78048 strains

To determine if alterations in in vivo gene expression could contribute to the virulence phenotypes observed with these strains and their *nrg1*ΔΔ mutants, we performed Nanostring analysis of ear tissue 24hr after infection with 78042, 78048 and their corresponding *nrg1*ΔΔ mutants (See Fig. S3A-D for volcano plots for each mutant compared to its parental strain; see Table SX for raw, processed and FC values along with FDR for each comparison). Compared to SN250 (15), the expression of some hypha-associated genes was reduced in 78042 and 78048 in vivo (Fig. 7A). Consistent with 78042 forming very few if any filaments, the expression of hypha-associated genes was reduced compared to the robustly filamentous 78048. In contrast, 78048 forms the same proportion of filaments as SN250 in vivo (Fig. 4, ref. 15) but the expression of hyphae-associated genes is significantly reduced, indicating that a similar level of hyphal morphogenesis does not result in similar levels of hyphae-associated gene expression in vivo. For example, the expression of candidalysin (*ECE1*), a key virulence-associated toxin (18), is 10-fold lower in 78048 compared to SN250 (Fig. 7A) but the strain has a similar median survival time and fungal burden as SN250 (Fig. 2D&H, ref. 14).

**Figure 7.**
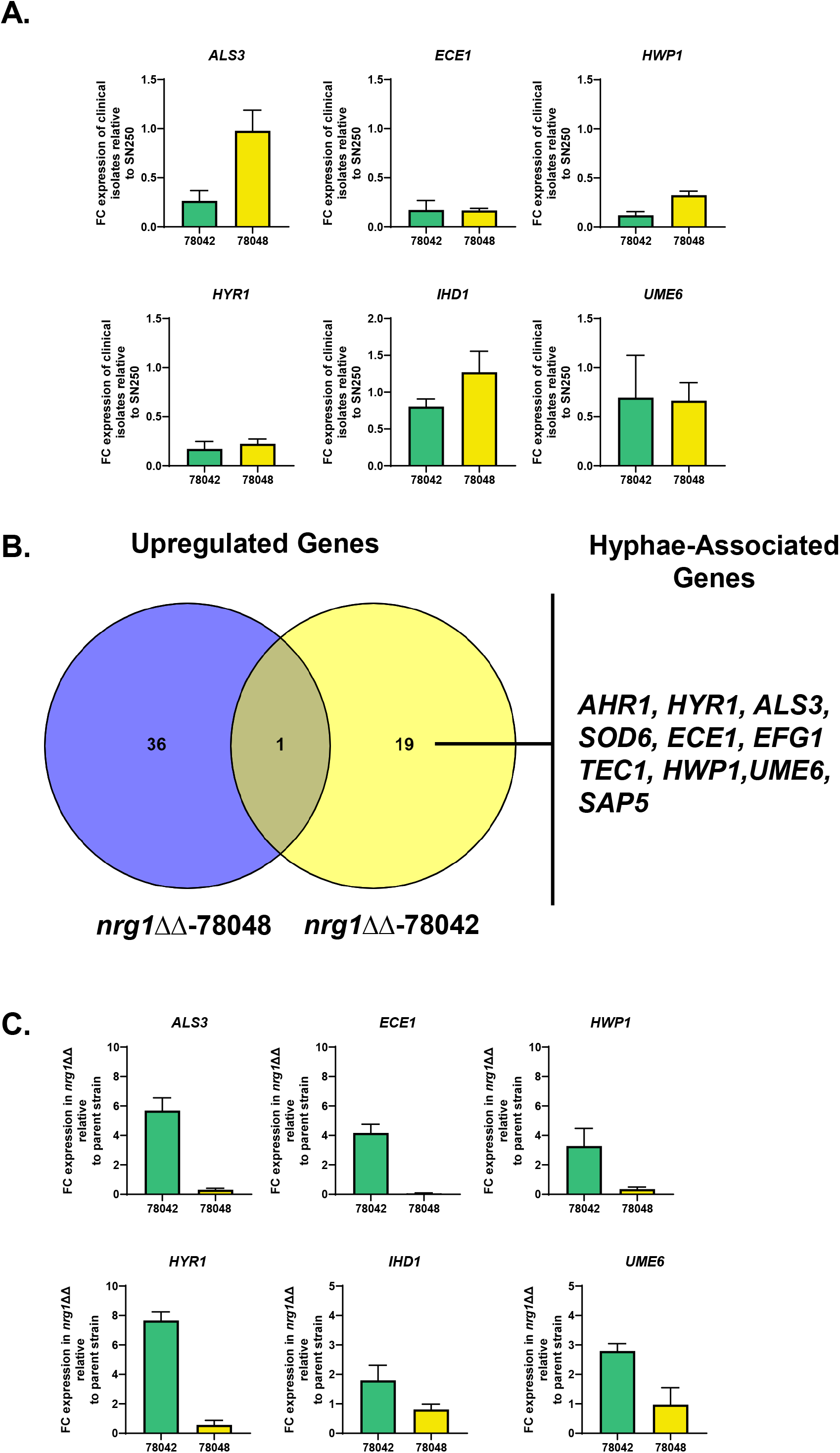
In vivo expression profiles of clinical isolates 78042 and 78048 as well as their respective *nrg1*ΔΔ mutants 24hr post-infection in ear. **A**. The expression for six hypha-associated genes in vivo in 78042 and 78048 compared to the strongly filamentous strain SN250. Bars indicate FC-relative to SN250 for three independent replicates. RNA was harvested as described in materials and methods 24hr post infection. * indicates statistically significant change relative to SN250 (±2-fold change in expression, FDR <0.1). **B**. Venn diagram of genes upregulated during infection in *nrg1*ΔΔ-78042 relative to 78042 and *nrg1*ΔΔ-78048 relative to 78048. See Tables S1&2 for FC data and significance data. **C**. Expression of the indicated hypha-associated genes in *nrg1*ΔΔ-78042 and *nrg1*ΔΔ-78048 normalized to expression in the parental strain. Bars indicate FC-relative in the *nrg1*ΔΔ mutant relative to the parental strains for three independent replicates. RNA was harvested as described in materials and methods 24hr post infection. * indicates statistically significant change relative to the parental change (±2-fold change in expression, FDR <0.1).

Next, we compared the in vivo expression profiles of the *nrg1*ΔΔ mutants to the 78042 and 78048 parental strains (See Fig. S4A&B for volcano plots, Table S2 for raw and processed data). As shown in the Venn diagram in Fig. 7B, the sets of upregulated genes in the two *nrg1*ΔΔ mutants were remarkably different. Surprisingly, only a single gene was upregulated in both *nrg1*ΔΔ-78042 and *nrg1*ΔΔ-78048 strains in vivo. Furthermore, the set of upregulated genes in the *nrg1*ΔΔ-78048 strain contained no hypha-associated genes while multiple key hypha-associated genes were upregulated in the *nrg1*ΔΔ-78042 strain including the positive transcriptional regulators of hyphae morphogenesis *AHR1*, *EFG1*, *TEC1*, and *UME6* (Fig. 7B, Table S2). The expression of selected hypha-associated genes in *nrg1*ΔΔ-78042 and *nrg1*ΔΔ-78048 are shown in Fig. 7C.

Notably, upregulation of these positive regulators of in vivo and in vitro morphogenesis does not lead to significant changes in the amount of in vivo filamentation observed in the *nrg1*ΔΔ-78042 mutant relative to its parental strain. Loss of *NRG1* does, however, trigger the increased expression of multiple hyphae-associated genes including *ECE1*, *HWP1*, and *HYR1* (Fig. 7C). Consistent with the upregulation of hypha-associated genes in the *nrg1*ΔΔ-78042 strain, the yeast-associated gene *YWP1* is downregulated 8-fold relative to 78042 (Table S2). Consequently, loss of *NRG1* in 78042 de-represses the expression of hypha-associated genes but does not lead to a corresponding increase in filamentation. The expression data further suggest that the increase in virulence observed for the *nrg1*ΔΔ-78042 strain relative to 78042 may be due to increased expression of hypha/virulence-associated genes such as *ECE1* or *HWP1*.

In contrast, deletion of *NRG1* in 78048 does not increase the expression of hypha-associated genes and, in fact, leads to the reduced expression of two canonical hypha-associated genes, *ECE1* and *ALS3* (Fig. 7C). Again, this observation fits with the reduced filamentation of the *nrg1*ΔΔ-78048 strain in vivo. *ECE1* expression is reduced 6-fold in 78048 compared to SN250 and is further reduced by 16-fold in the *nrg1*ΔΔ-78048 mutant. Thus, it seems likely that the dramatically reduced expression of *ECE1* in the *nrg1*ΔΔ-78048 mutant may also contribute to its reduced virulence (19), although reduced filamentation or reduced expression of other genes could also play a role.

## Discussion

Using a set of clinical isolates with relatively poor in vitro filamentation, we explored the relationship between in vitro and in vivo filamentation and virulence. From this analysis, we can draw three conclusions. First, in vitro filamentation phenotypes using standard conditions, including host like conditions, correlate with neither infectivity nor virulence in the four isolates that we examined. This supports the conclusion of a previous study using historical virulence data (4, 7). Second, the two isolates (94015 and 78042) that do not filament in vivo are less virulent than the two strains (57055 and 78048) that do filament in vivo. These findings suggest that at least some of the discordance between in vitro filamentation phenotypes and in vivo virulence is due to the fact that the infection environment drives filamentation in some strains that do not undergo filamentation well in vitro. Third, *C. albicans* strains that are unable to filament in vitro or in vivo (94015 and 78042) are still able to establish infection in the kidney, the major target organ of this model. This is consistent with previous observations that yeast-locked mutants of SC5314 also establish kidney infection (13).

At the outset of the project, we somewhat naively hypothesized that deletion of *NRG1* would increase the filamentation of these poorly filamenting strains and potentially increase their virulence as well. In vitro, deletion of *NRG1* leads to a consistent phenotype in which the strains show increased pseudohyphae formation and increased expression of hypha-associated genes. Although there are some strain specific distinctions in the DEGs sets, these are generally minor with the exception of *nrg1*ΔΔ-94015 which lacks Efg1. Thus, at a broad level, the function of Nrg1 is consistent across multiple clinical strains.

There are, however, phenotypic differences between *nrg1*ΔΔ-mutants in the clinical isolates and the SN250-derived mutant. First, as previously described by many others (10, 11), deletion of *NRG1* in SC5314-derivatives leads to constitutive pseudohyphae formation in the absence of hypha-inducing stimuli whereas the yeast morphology is predominant in *nrg1ΔΔ* mutants generated in the four poorly filamenting strains. However, this phenotype is one of degree rather than a simple binary state because all four poorly filamenting do form pseudohyphae in non-inducing conditions. Thus, additional factors must be present in SN250 that that mediate the strong constitutive pseudohyphal phenotype; the poor filamenting strains apparently lack these factors. Second, upon in vitro induction, the *nrg1*ΔΔ-SN250 forms predominately hyphae while the clinical isolate-derived *nrg1*ΔΔ mutants generate a mixture of yeast and pseudohyphae with almost no true hyphae observed. This further suggests that the mechanisms which make SC5314, and its derivatives robustly filamentous and that are lacking in the clinical isolates, operate downstream of the relief of Nrg1 repression step.

It is unclear what genes or pathways drive the robust filamentation of SC5314, although we have recently found that a gain-of-function SNP present in the transcription factor Rob1 contributes to the strong filamentation of SC5314; we have not identified this SNP in any other sequenced strain (14); deletion of *NRG1* in a SC5314-derived mutant with only the more prevalent *ROB1* allele did not block hyphae formation (Wakade and Krysan, unpublished), indicating that this cannot explain the differential effect of *NRG1* deletion between these strains and SC5314. On the other hand, it is still possible that strain-specific alterations in the function of positive regulators of filamentation may be present in these strains. However, an examination of the mutations in 78042 and 78048 (relative to SC5314) did not reveal any obvious alterations in key hypha-related regulators (4). Thus, additional studies will be required to identify the hypha-associated pathways that are lacking in these strains. The increasing availability of diverse *C. albicans* clinical isolates should facilitate that search.

Whereas the in vitro phenotypes and expression profiles of the four clinical isolate-derived *nrg1*ΔΔ mutants are fairly consistent, they show phenotypic heterogeneity in vivo. We initially hypothesized that deletion of *NRG1* would increase virulence in general. However, this hypothesis was confirmed conclusively for only one of the four strains. Strikingly, the increased virulence of the *nrg1*ΔΔ-78042 strain was not due to increased filamentation but instead more likely to be due to increased expression of hypha-associated genes such as *ECE1* in cells that remained in yeast morphology in vivo. Contrary to our hypothesis, deletion of *NRG1* lead to reduced virulence in the two most virulent of our four strains (57055 and 78048). This difference was not due to a failure to establish initial infection or due to clearance of the mutants from the kidneys.

In the case of the *nrg1*ΔΔ-78048 mutant, the reduced virulence is likely to be due to multiple factors including altered extent of filamentation and filament length. Most notable, however, is the reduced expression of the toxin *ECE1* in the *nrg1*ΔΔ-78048 mutant relative to the parental strain. In vitro and in vivo, deletion of *NRG1* increases the expression of *ECE1* in multiple strain background. In vitro, the *nrg1*ΔΔ-78048 mutant has increased expression of *ECE1* but in vivo it is reduced by 16-fold. These results indicate that, under some conditions and in some strains, the loss of the repressor Nrg1 leads to decreased expression of some its canonical targets. These seemingly paradoxical findings are consistent with a similar strain dependent effect of the *NRG1* deletion reported by the Mitchell lab and highlight the complexity and plasticity of transcriptional networks across different *C. albicans* strains (6).

Our characterization of the in vivo filamentation and virulence phenotypes of the four parental strains also provides some explanations for the discordance between in vitro filamentation and virulence. Specifically, the ability of some strains to filament in vivo but not in vitro are likely to explain some of the phenotypic contradictions. Since some strains that cannot filament in vivo are less virulent than those that do, we propose that filamentation remains a general virulence trait for *C. albicans*. However, our data also suggest that alteration in the expression of hypha/virulence associated genes in the absence of strong changes in filamentation is also likely to contribute. It is important to consider that even the strain lacking a functional allele of the master virulence and filamentation regulator Efg1 caused disease in these models and, since it was isolated from the blood of a patient, in humans to some extent. As such, we cannot forget the role of the host and the reasons or risk factors that predisposed them to invasive candidiasis.

## Materials and methods

### Strains, cultivation conditions and media

The *C. albicans* clinical isolate strains (94015, 57055, 78042, and 78048) used in this study was obtained from the Soll laboratory (ref. 7). Standard recipes were used to prepare yeast peptone dextrose (YPD) (20) and all *C. albicans* strains were precultured overnight in YPD medium at 30^0^C. RPMI medium was purchased and supplemented with bovine serum (10%, vol/vol). For *in vitro* hyphal induction, *C. albicans* strains were incubated overnight at 30^0^C in YPD media, harvested, and diluted into RPMI + 10% serum at a 1:50 ratio and incubated at 37^0^C for 4 h (9).

### Strain construction

*C. albicans* transformation was performed using the standard lithium acetate transformation method (21). The homozygous mutant strains of *C. albicans* were constructed using the transient CRISPR/Cas9 method (21). Oligonucleotides and plasmids used to generate the mutant strains in this study are listed in Table S3. The *NAT1-Clox* system (22) was used to generate the *NRG1* KO strains. Briefly, both copies of the *NRG1* was deleted by amplifying *NAT1-Clox* cassette from *NAT1-Clox* plasmid with primer pairs *NRG1.P1* and *NRG1.P2*, and by using sgRNA targeting both alleles of *NRG1* gene. The resultant transformants were selected on YPD containing 2.5 mM methionine and 2.5 mM cysteine along with 200 µg/ml nourseothricin (Werner Bioagents, Jena, Germany). The correct transformants were confirmed by standard PCR methods with primer pairs *NRG1.P5* and *NRG1.P6*. Oligonucleotide sequences are provided in Table S3.

Fluorescently labeled strains were generated by using p*ENO1-NEON-NAT1* and p*ENO1-iRFP-NAT1* plasmids as previously described (9) and the resultant transformants were selected on YPD containing 200 µg/ml nourseothricin (Werner Bioagents, Jena, Germany). The reference strain was tagged with green fluorescent protein (NEON) whereas the mutant strains were tagged with iRFP.

### In vitro characterization of *C. albicans* morphology

Induced cells were fixed with 1% (v/v) formaldehyde. Fixed cells were then imaged using the Echo Rebel upright microscope with a 60x objective. The assays were conducted in triplicates on different days to confirm reproducibility.

### Mouse model of disseminated candidiasis

5 to 6 weeks old, female CD1 mice (Envigo) were inoculated with 1 X 10^6^ CFU (200 µl) of reference or mutant strain by lateral tail vein injection. Group of five mice were used to assess the kidney fungal burden. Mice were monitored daily and were sacrificed three days after infection as per the IACUC guidelines to determine the kidney fungal burden. Two kidneys were homogenized together in 1 ml sterile PBS and 10 µl of 10-fold serial dilutions of kidney extracts were spotted on to the YPD plates and fungal burden was determined with the Unpaired t test.

To determine the virulence of the reference or mutant strains, 10 female CD1 mice per group were inoculated with the 1 X 10^6^ CFU (200 µl) by lateral tail vein injection. Mice were monitored daily and when a mouse exhibited any signs of extreme morbidity such as hunched back, head tilt or tremors, mouse was euthanized as per the IACUC guidelines. Survival curves were plotted on a Kaplan-Meier curve, and Log-rank (Mantel-Cox) test was used to determine the statistical difference of the curves (Graph Pad Prism).

### In vitro and in vivo RNA Extraction

RNA extractions for the *in vitro* and *in vivo* samples were carried our as described previously (15). Briefly, for *in vitro* RNA extractions, three independent samples were grown ON in YPD at 30^0^C, washed twice in the PBS and diluted at 1:50 ratios in the RPMI + 10% serum and incubated at 37^0^C for 4 hours. Cells were collected, centrifuged for 2 min at 11K rpm at room temperature and RNA was extracted as per the manufacturer protocol (MasterPure Yeast RNA Purification Kit, Cat. No. MPY03199). Extraction of RNA from mouse ear was carried out as described previously (15). Briefly, 24 hours post infection, mouse was euthanized following the protocol approved by the University of Iowa IACUC. The *C. albicans* injected mouse ear was excised and placed in an ice-cold RNA later solution. Subsequently, the ear was transferred to a mortar, flash-frozen using liquid nitrogen and ground to the fine powder. The powder was collected into 5 ml centrifuge tube and 1 ml of ice-cold Trizol was added. The samples were placed on a rocker at RT for 15 min and then centrifuged at 10K rpm for 10 min at 4^0^C. The cleared Trizol was then collected without dislodging the pellet into 1.5 ml Eppendorf tube and 200 µl of RNase free chloroform was added. The tubes were shaken vigorously for 10-15s and kept at RT for 5 min. Further the samples were centrifuged at 12K for 15 min at 4^0^C. The cleared aqueous layer was then collected to a new 1.5 ml RNAase free Eppendorf tube and RNA was further extracted using the Qiagen RNeasy kit protocol.

### NanoString analysis of gene expression in vitro and in vivo

NanoString analysis was carried out as described previously (15). Briefly, in total, 80 ng of *in vitro* or 400 ng of *in vivo* RNA was added to a NanoString codeset mix and incubated at 65^0^ C for 18 hours. After hybridization reaction, samples were proceeded to nCounter prep station and samples were scanned on an nCounter digital analyzer. nCounter .RCC files for each sample were imported into nSolver software to determine the quality control metrics. Using the negative control probes the background values was defined and used as a background threshold and this value is subtracted from the raw counts. The resulting background subtracted total raw RNA counts were first normalized against the highest total counts from the biological triplicates and then to the highest total counts for the samples. The statistical significance of changes in gene expression was determined using Benjamini-Hochberg method at an FDR of 0.1. The raw, normalized and analyzed expression data are provided in the Tables S1 and S2.

### In vivo imaging and filamentation scoring

The inoculation and imaging of mice ear were carried out as described previously (9, 15). Acquired multiple Z stacks (minimum 15) were used to score the yeast vs filamentous ratio. The cells were considered as a “Yeast” if the cells were round and/or budded cells. Furthermore, yeast cells were required not to project through multiple Z stacks. The cells were considered as a “filamentous” if the cells contain intact mother and filamentous which was at least twice the length of the mother body. A minimum of 100 cells from multiple fields were scored. Paired Student’s *t* test with Welch’s correction (*P*>0.05) was used to define the statistical significance which was carried out using GraphPad prism software. Filament length of the *in vivo* samples were measured as described previously (15, 23). Briefly, a Z stacks image of the reference or mutant strain were opened in an ImageJ software and the distance between mother neck to the tip of the filament was measured. At least 50 cells per each strain from multiple fields were measured. Statistical significance was determined by Mann-Whitney U test (*P*>0.05).

## Acknowledgements

This work was supported by NIH grant R01AI133409 (DJK). We thank Aaron Mitchell (Georgia) for communicating results prior to publication. We also thank Scott Filler (UCLA) for suggestions and discussion.

## Author Contributions

Conceptualization: Damian J. Krysan

Formal analysis: Rohan S. Wakade, Melanie Wellington, and Damian J. Krysan

Investigation: Rohan S. Wakade

Methodology: Rohan S. Wakade, Melanie

Wellington Supervision: Melanie Wellington, Damian J. Krysan

Writing-original draft: Damian J. Krysan

Writing-review and editing: Damian J. Krysan, Rohan S. Wakade

## Supplementary Material Legends

**Fig. S1.**
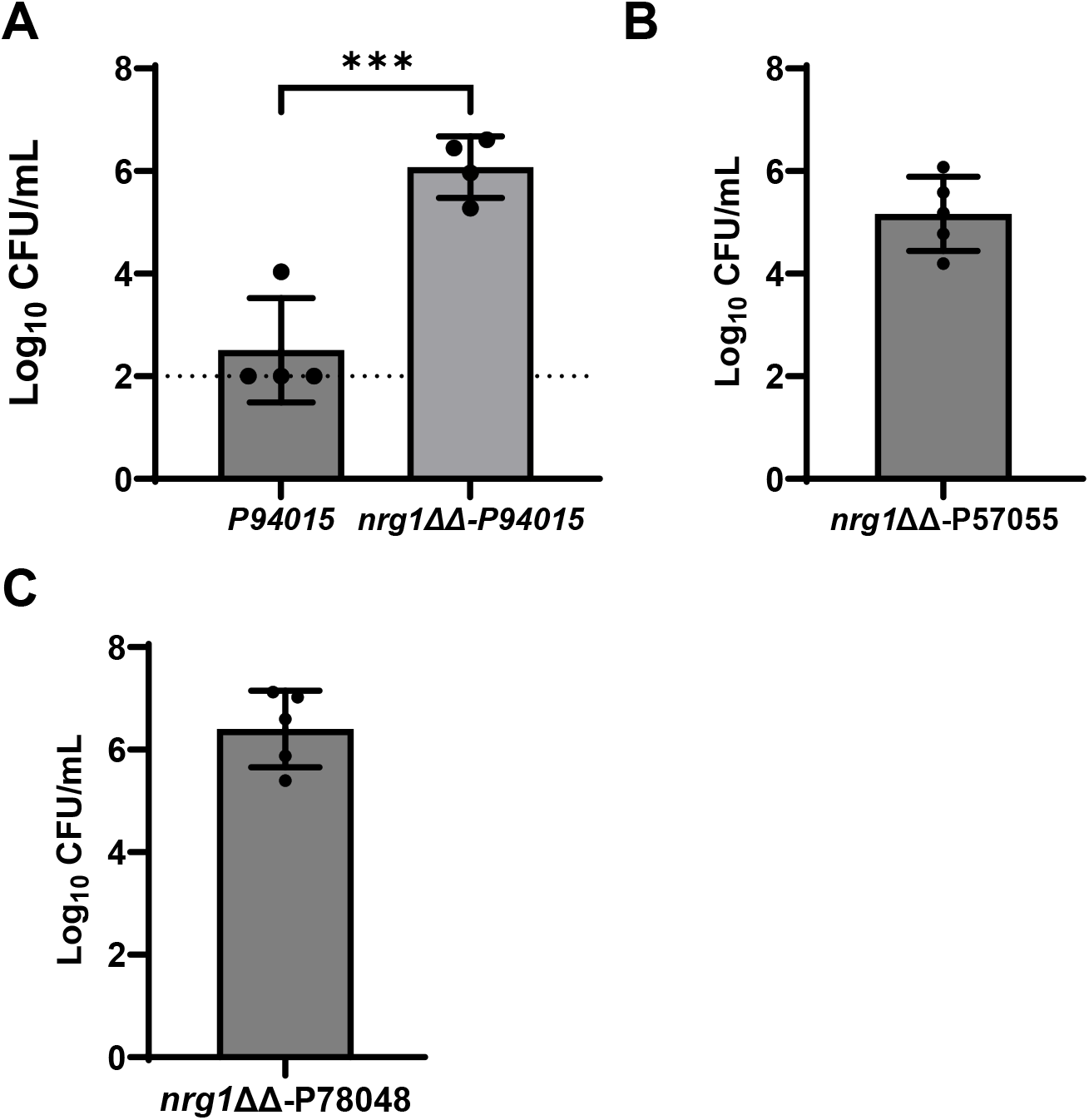
Kidney fungal burden for surviving animals in virulence experiments. Kidney fungal burden from animals surviving at the end of virulence assays (**Fig. 2**). **A**. 94015. **B**. *nrg1*ΔΔ-57055. **C**. *nrg1*ΔΔ-78048. Bars indicate mean of log^10^ fungal burden (CFU/mL) with individual mice shown as points and error bars indicating standard deviation. Differences between groups were analyzed by Student’s t test (NS: p> 0.05 and *** p< 0.0001).

**Fig. S2.**
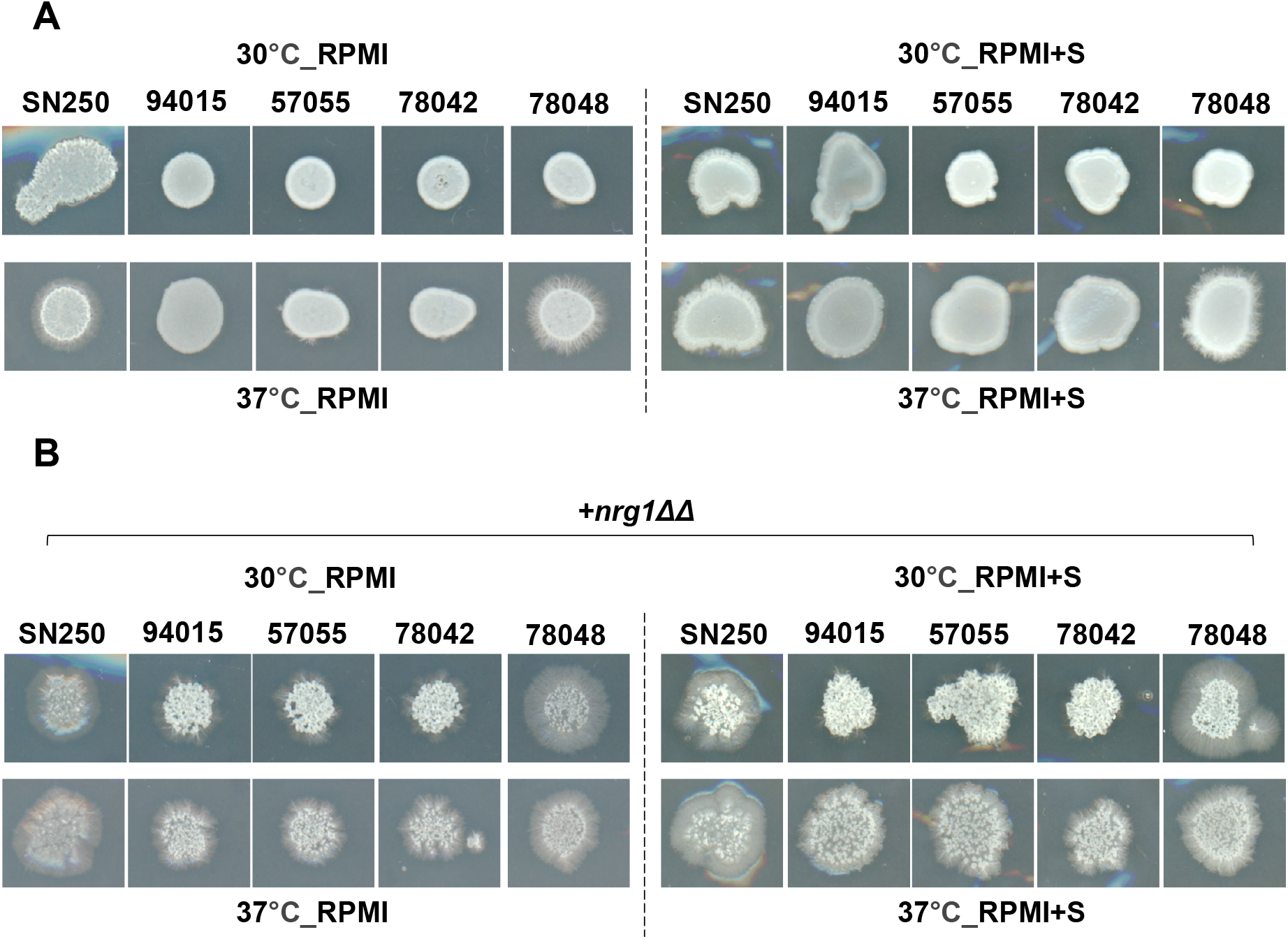
Filamentation phenotypes of clinical isolates and their corresponding *nrg1*ΔΔ mutants. The indicated clinical isolates (A) and corresponding *nrg1*ΔΔ mutants were plated on RPMI and RPMI+10% bovine calf serum and incubated at either 30°C or 37°C for 3 days and photographed. The experiments were performed in biological duplicate and phenotypes were similar in both experiments.

**Fig. S3.**
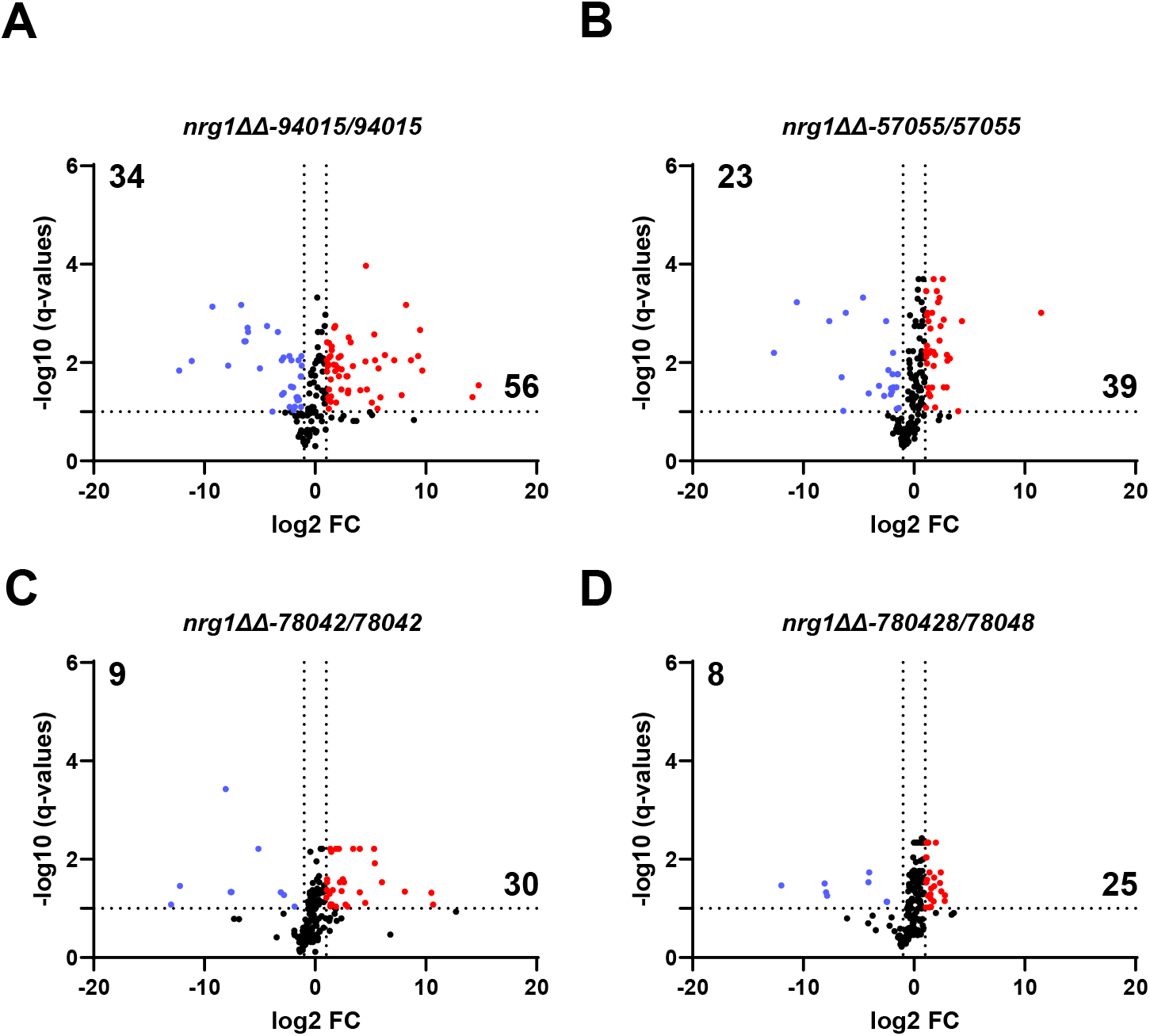
Volcano plots comparing the in vitro expression profiles for clinical isolates to their corresponding *nrg1*ΔΔ mutants. Volcano plots of gene expression of a set of 185 genes as characterized by Nanostring nCounter. The expression was normalized to the strongly filamenting reference strain SN250. The horizontal bar indicates FDR 0.1 (Benjamini-Hochberg) and the vertical line indicates log^2^ = 1 of the fold change which are the cutoff values for the definition of differentially expressed genes. **A.** 94015; **B.** 57055. **C.** 78042. **D.** 78084.

**Fig. S4.**
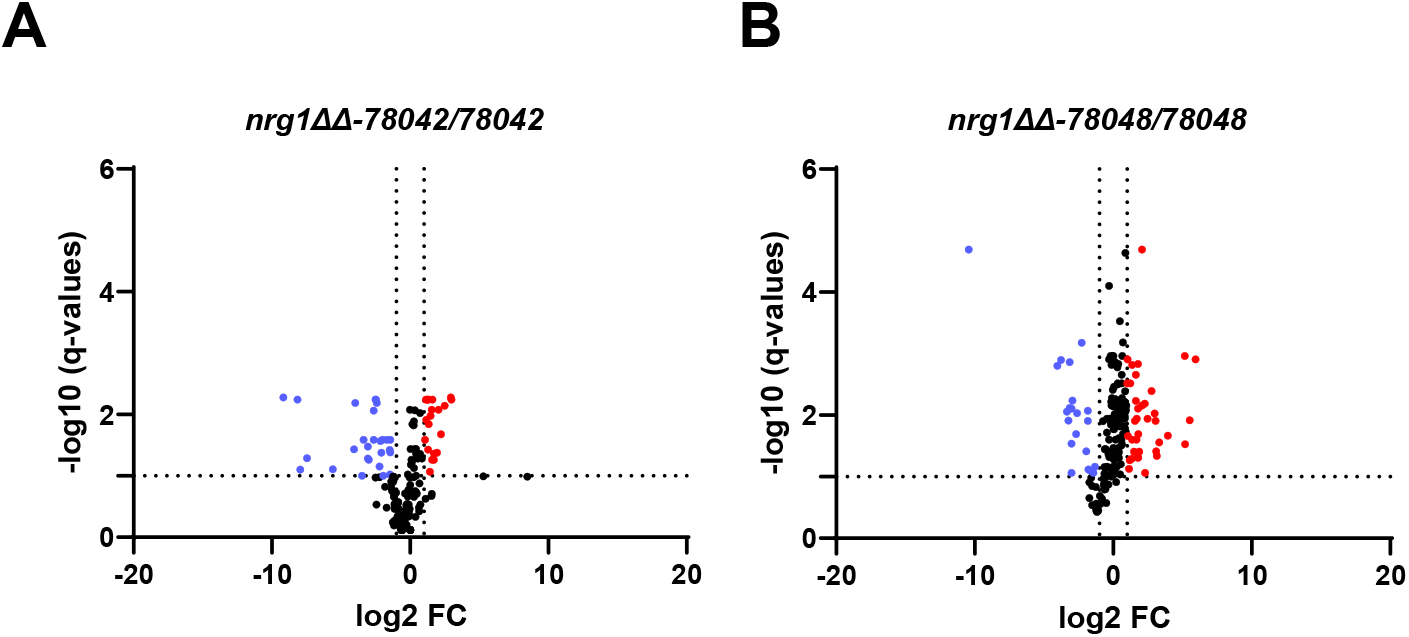
Volcano plots comparing the in vivo expression profiles for clinical isolates 78042 and 78048 to their corresponding *nrg1*ΔΔ mutants. Volcano plots of gene expression of a set of 185 genes as characterized by Nanostring nCounter. The expression was normalized to the strongly filamenting reference strain SN250. The horizontal bar indicates FDR 0.1 (Benjamini-Hochberg) and the vertical line indicates log^2^ = 1 of the fold change which are the cutoff values for the definition of differentially expressed genes. **A.** 78042. **B.** 78084.

**Table S1. Nanostring expression data for in vitro experiments**. RNA was harvested from the indicated strains 4hr after induction with RPMI+10%BCS. Raw mRNA counts, normalized counts, average counts, fold-change from reference strain, Student t test, and Benjamini-Hochberg adjusted FDR are provided for each gene and condition (two to three biological replicates). Differentially expressed genes were defined as ±2-fold-change in expression with FDR<0.1. Fold-change values shown in red are significantly downregulated and those shown in green are significantly upregulated. Summaries of all differentially expressed genes are also provided.

**Table S2. Nanostring expression data for in vivo experiments**. RNA was isolated from mouse ear tissue 24hr post-infection. Raw mRNA counts, normalized counts, average counts, fold-change from reference strain, Student t test, and Benjamini-Hochberg adjusted FDR are provided for each gene and condition (two to three biological replicates). Differentially expressed genes were defined as ±2-fold change in expression with FDR<0.1. Fold-change values shown in red are significantly downregulated and those shown in green are significantly upregulated. Summaries of all differentially expressed genes are also provided.

**Table S3. Oligonucleotides used for strain construction and analysis.**

